# Healthy aging interventions reduce non-coding repetitive element transcripts

**DOI:** 10.1101/2020.06.25.172023

**Authors:** Devin Wahl, Alyssa N. Cavalier, Meghan Smith, Douglas R. Seals, Thomas J. LaRocca

## Abstract

Transcripts from non-coding repetitive elements (RE) in the genome may be involved in aging. However, they are often ignored in transcriptome studies on healthspan and lifespan, and their role in healthy aging interventions has not been characterized. Here, we analyze RE in RNA-seq datasets from mice subjected to robust healthspan- and lifespan-increasing interventions including calorie restriction, rapamycin, acarbose, 17-α-estradiol, and Protandim. We also examine RE transcripts in long-lived transgenic mice, and in mice subjected to high-fat diet, and we use RNA-seq to investigate the influence of aerobic exercise on RE transcripts with aging in humans. We find that: 1) healthy aging interventions/behaviors globally reduce RE transcripts, whereas aging and age-accelerating treatments increase RE expression; and 2) reduced RE expression with healthy aging interventions is associated with biological/physiological processes mechanistically linked with aging. Thus, RE transcript dysregulation and suppression are likely novel mechanisms underlying aging and healthy aging interventions, respectively.

## INTRODUCTION

Older age is the greatest risk factor for the development of most chronic diseases (1). Accordingly, recent large-scale ‘omics’ studies have aimed to characterize novel genes and biological pathways that influence aging, and to identify related interventions (e.g., pharmaceutical compounds, exercise, nutrition) that increase longevity and healthspan (2, 3). Indeed, advances in transcriptomics (e.g., RNA-seq) have led to important insight on many genes and pathways linked with ‘the hallmarks of aging’ and broader health outcomes (4). However, most of these studies have focused on coding sequences—a small fraction of the genome. Non-coding, repetitive elements (RE, >60% of the genome) have been particularly neglected as ‘junk DNA’ (5), despite growing evidence that they have many important biological functions (6).

RE include DNA transposons, retrotransposons, tandem repeats, satellites and terminal repeats (7). A major fraction of RE, mainly DNA transposons and retrotransposons (e.g., LINEs, SINEs, LTRs), are transposable elements (TE) with the ability to propagate, multiply and change genomic position (6). Most RE/TE are in genomic regions that are chromatinized and suppressed (inactive), but recent reports show that certain TE become active during aging, perhaps due to reduced chromatin architecture/stability (e.g., histone dysregulation) (8). Activation of these specific TE may contribute to aging by causing genomic and/or cellular damage/stress (e.g., inflammation) (9). However, we recently reported that aging is associated with a progressive, *global* increase in transcripts from most RE (not only TE) in model organisms and humans (10). This global dysregulation of RE may have an important, more general role in aging, as RE transcripts have been linked with other key hallmarks of aging including oxidative stress and cellular senescence (11). In fact, it has been suggested that RE dysregulation itself may be an important hallmark of aging (12). If so, a logical prediction would be that interventions that increase health/lifespan and reduce hallmarks of aging (e.g., calorie restriction [CR], select pharmacological agents and exercise) should also suppress RE/TE. Limited evidence suggests this may be true for CR and certain TE in *Drosophila* (13), but global RE/TE expression has not been well studied in this context.

## RESULTS AND DISCUSSION

To determine if global RE transcript suppression might be a mechanism underlying healthy aging interventions, we first analyzed RE in an RNA-seq dataset on livers from young and old mice and old mice subjected to life-long (24 months) CR (14). We found a small, but significant age-related increase in most major RE transcript types in this dataset, consistent with our previous work (15) and others’ (16). However, this effect was significantly attenuated with CR (**Fig. 1A**). Based on this novel evidence of RE suppression by CR (arguably the strongest health/lifespan-enhancing intervention), we looked to confirm our results in an additional, large dataset including RNA-seq on livers from mice subjected to different durations of CR and pharmacological interventions known to increase health/lifespan (rapamycin, acarbose, 17-α-estradiol [17aE], and Protandim), as well as data on transgenic, long-lived mice (2) (**Fig. 1 and Supplementary Data**). We found that long-term (8 months) CR caused a significant, global reduction in RE transcripts (**Fig. 1B**). Furthermore, we found that both long-term (8 months) rapamycin and acarbose treatments were associated with a comparable, broad reduction in RE transcripts (**Fig. 1B**), consistent with the notion that these compounds are ‘calorie restriction mimetics’ and may act via similar pathways (17). This effect was particularly clear when we examined RE/TE reductions by major sub-type (**Fig. 1C**). Short-term (2 months) interventions with other healthy aging compounds influenced RE transcript levels to various degrees, although reductions were more pronounced with CR and Protandim, which is thought to activate endogenous antioxidant defenses (**Fig. 1D**). Interestingly, the authors of the original study (2) observed similar variability in gene expression patterns, suggesting time/treatment-specific transcriptome effects. We also found a significant influence of growth hormone receptor knockout (GHRKO, a transgenic longevity model) on the main RE transcript types (**Fig. 1E**). Moreover, in a separate dataset (18), we found that high-fat diet (HFD, a common ‘pro-aging’ intervention) significantly *increased* all major RE/TE (**Fig. 1F**). Collectively, these results support the idea that global RE transcript levels are linked with healthspan/lifespan, as they are reduced by most “gold standard” anti-aging interventions and increased by adverse, pro-aging treatments.

**Figure 1.**
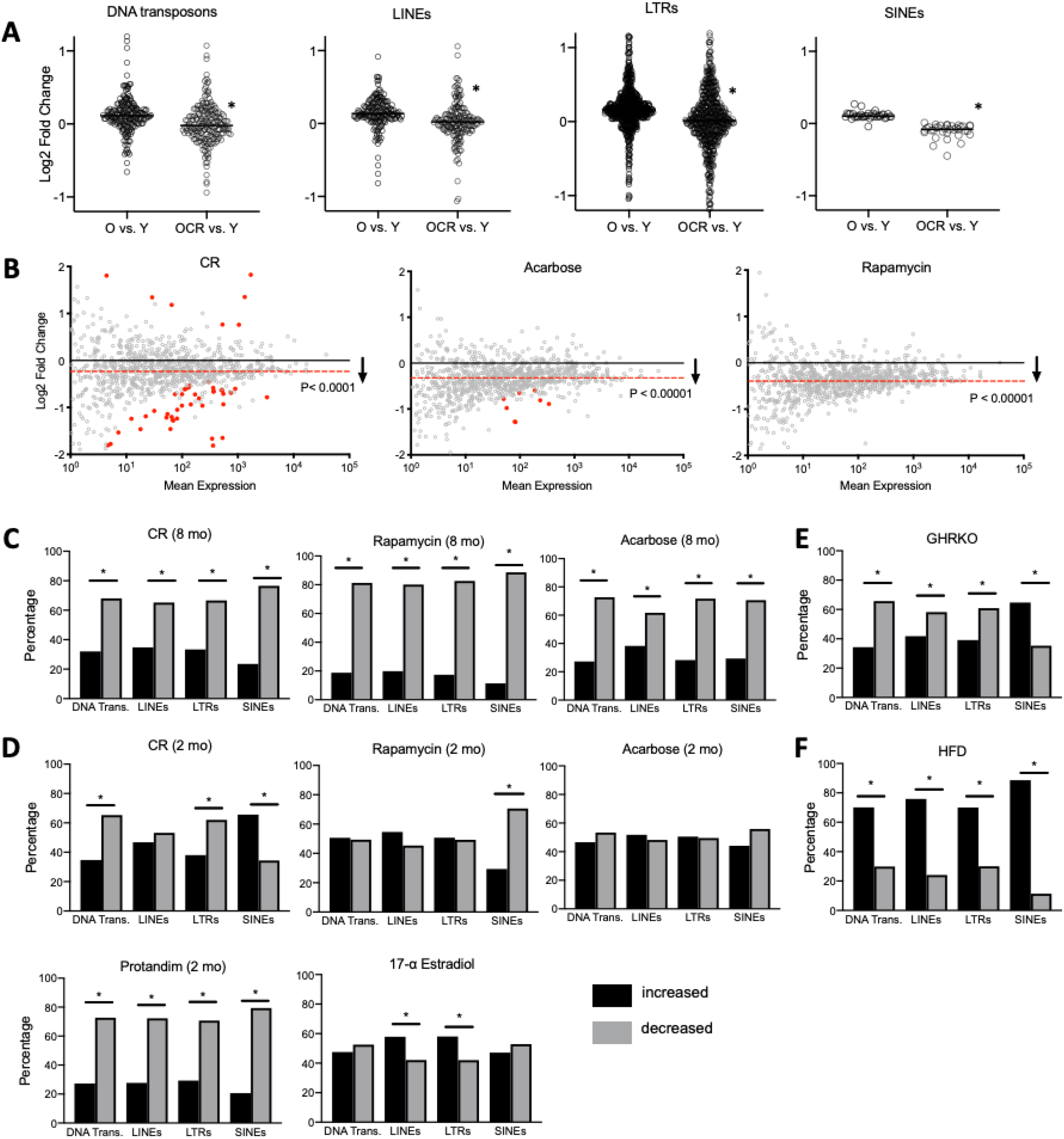
Healthy aging interventions reduce RE transcripts. **(A)** Age-related increases in the major types of RE transcripts in old (O) or old calorie-restricted (OCR) vs. young (Y) mice. N=3/group, *p<0.0001, two-tailed t-test. All individual RE shown. **(B)** MA plots showing significant decreases in global RE transcripts with long-term (8-month) healthy aging interventions. Significantly reduced or increased RE (FDR<0.1, identified using Deseq2) shown in red, and average transcript reduction indicated by red line. Likelihood of increased/decreased distribution calculated by chi-square analysis. **(C-F)** Percentage of RE transcripts by type increased or decreased with long- or short-term (2-month) interventions (*p<0.05, chi-square analysis). All relevant data/samples from datasets GSE92486, GSE131901 and GSE87565 were used for analyses (N=3-6 mice/group), and raw data are provided in the supplementary data file.

Next, we examined similarities in the effects of healthy aging interventions on RE by sub-type/family (**Fig. 2 and Supplementary Data**). Again, we observed variable patterns of RE family expression with short-term treatments (**Fig. 2A**). With long-term treatment, CR and rapamycin influenced RE/TE families most similarly, and most transcripts were decreased with all treatments (**Fig. 2B**), and in GHRKO mice (**Fig. 2C**). We next determined which specific RE transcripts were commonly decreased/increased among all treatments. Despite the variable RE/family expression patterns noted above, short-term treatments modulated many of the same transcripts (**Fig. 2D and Supplementary Data**). Long-term treatments also decreased/increased a large number of common transcripts (518 and 92, respectively) (**Fig. 2E and Supplementary Data**). Consistent with the idea that global RE modulation is linked with healthy aging interventions, we did not notice any particular enrichment for specific RE/TE types in these common transcripts. However, we did note that endogenous retrovirus (ERV) RE transcripts were the most decreased with all long-term treatments and in GHRKO mice. Interestingly, ERVs have been implicated in aging and several diseases of aging including neurodegenerative disorders (19, 20), suggesting that these RE could be an important therapeutic target.

**Figure 2.**
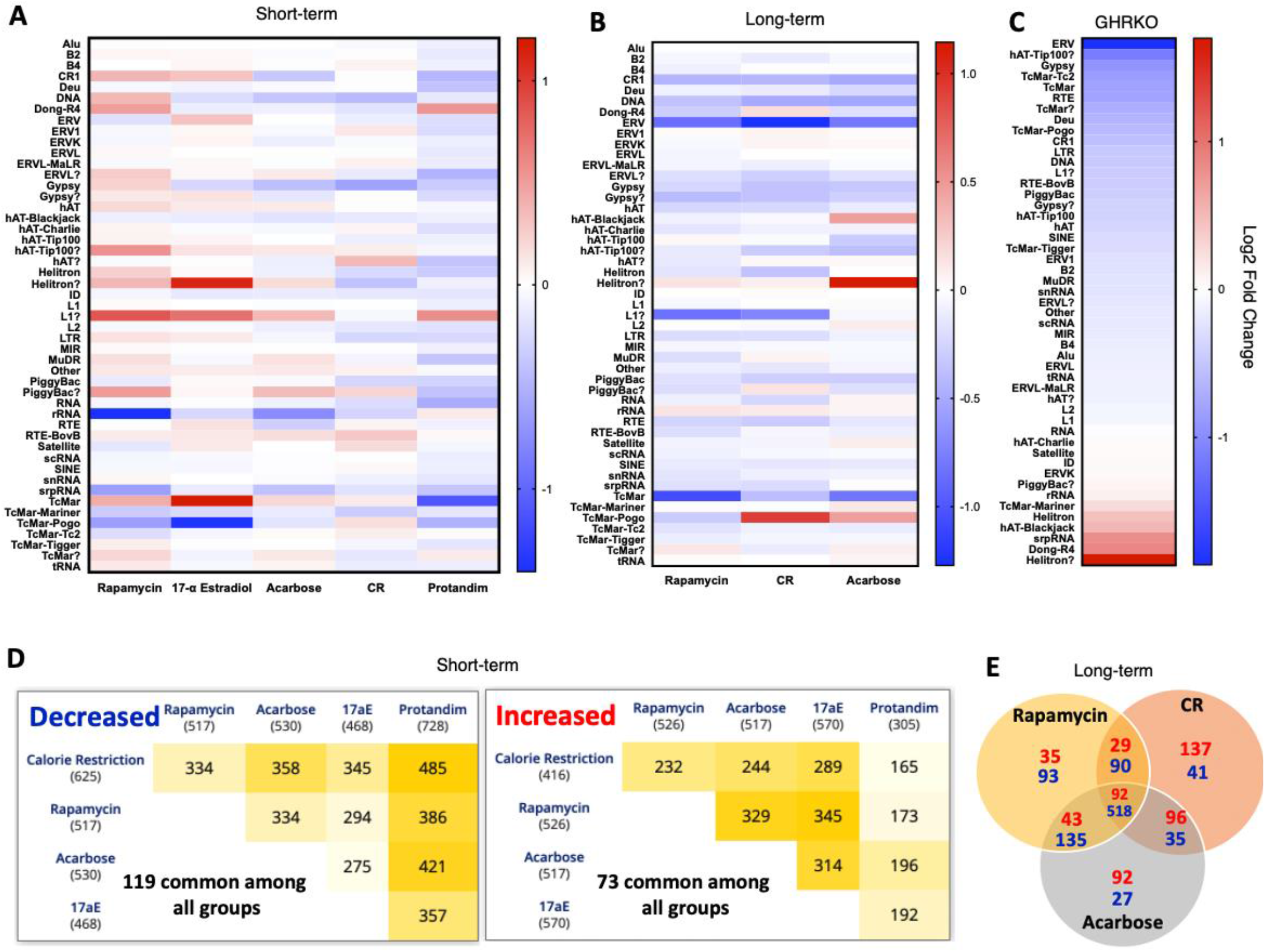
Healthy aging interventions reduce most RE families and many similar, individual RE transcripts. **(A-C)** Heatmaps showing RE transcript families increased (red) and decreased (blue) by short-term interventions, long-term interventions, and in GHRKO mice. **(D)** Pairwise intersections showing the number of common decreased or increased RE transcripts with short-term interventions. **(E)** Venn diagrams showing the number of common decreased (blue) or increased (red) RE transcripts with long-term interventions. All relevant data/samples from dataset GSE131901 were used for analyses (N=3-6 mice/group), and raw data are provided in the supplementary data file.

Identifying potentially targetable biological mechanisms linking reduced RE expression with healthy aging interventions will require future experiments. However, to provide initial insight, we examined correlations among gene and RE expression patterns in mice subjected to long-term CR, rapamycin and acarbose (treatments that reduced RE transcripts the most). To do this, we conducted a weighted gene correlation network analysis (WGCNA) on both gene and RE transcript counts (**Fig. 3 and Supplementary Data**). Although gene/RE signatures across interventions were not strikingly similar, we identified one WGCNA module (green) that decreased significantly with all interventions (**Fig. 3A**). This module contained numerous DNA transposons and several LINE, ERV and LTR transcripts. A gene ontology (GO) analysis of the module also showed significant enrichment for many biological processes, including several linked with aging and disease (**Fig. 3B**). In fact, the most specific GO terms included protein deacetylation, DNA repair and immune activation/response pathways. The other gene/RE modules that decreased with CR and/or rapamycin were also enriched for specific GO terms including DNA repair, DNA/RNA processing, histone modifications and stress responses (**Fig. 3B**). These exploratory analyses do not definitively link reduced RE expression with such processes, but they are consistent with current thinking that age-related RE transcript accumulation could cause DNA damage (21) and immune activation/inflammation (22), and that RE dysregulation may be due to age-associated changes in chromatin/histones (23).

**Figure 3.**
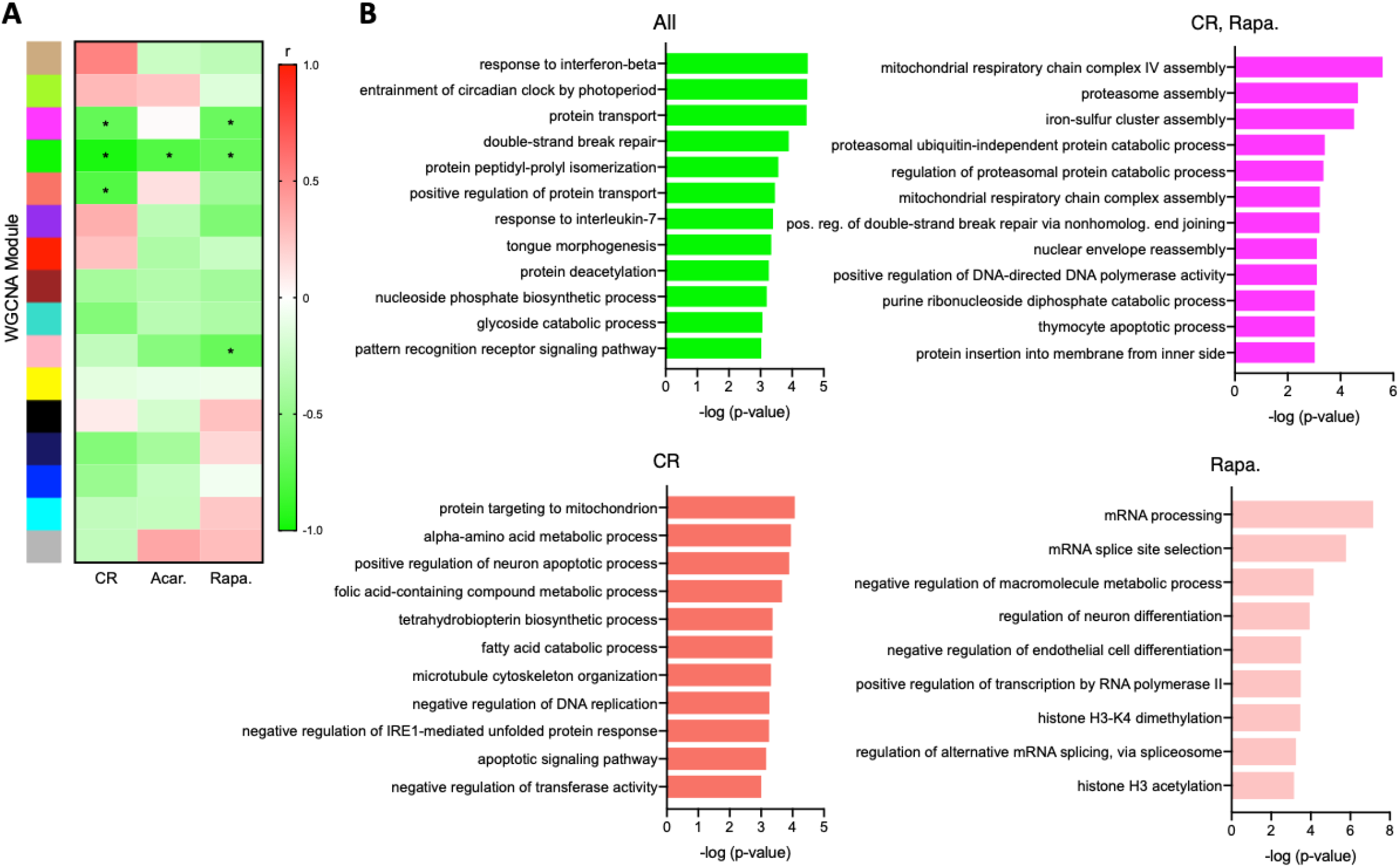
Gene expression patterns associated with RE transcript reductions. **(A)** WGCNA analysis heatmap of gene/RE modules influenced by long-term interventions (*modules that changed significantly with interventions, P<0.001 in WGCNA). **(B)** Most specific biological processes (gene ontology terms) in each significant WGCNA module. Exact p-values are noted in the supplementary data file, and all available samples/data from dataset GSE131901 were used for analyses (N=6 mice/group).

There is little or no RNA-seq data on true long-term CR or healthy aging compounds in older humans, as these are challenging clinical interventions to conduct (24). However, one well-studied intervention/behavior associated with increased healthspan and biological effects similar to CR is aerobic exercise (25, 26). Others have studied the transcriptomic effects of exercise interventions on select tissues (27), but proportionally, no studies have been nearly as long as an 8-month mouse intervention (~25% of the animal’s life). Therefore, as initial proof of concept, we conducted a cross-sectional study to determine if long-term exposure to this healthy aging behavior has the potential to reduce RE transcripts (**Fig. 4 and Supplementary Data**). We performed RNA-seq and gene/RE expression analyses on peripheral blood mononuclear cells (PBMC) from: 1) young and older sedentary adults; and 2) older habitually (≥5 years) exercising adults (**Supplementary Table**). Consistent with other reports (28), we found that older age was associated with altered PBMC gene expression, but these changes were largely attenuated in exercising older adults (**Fig. 4A**). Moreover, in support of the idea that healthy aging interventions/behaviors may reduce RE expression in humans, we observed a clear increase in global RE transcript levels in older sedentary adult PBMC, but this effect was strongly attenuated with exercise (**Fig. 4B**). We also found that maximal aerobic exercise capacity (VO_2_ max) was inversely related to a composite count of RE that are significantly increased with aging (**Fig. 4C**), suggesting that greater aerobic fitness (and perhaps exposure to aerobic exercise) is directly linked with reduced RE expression. Interestingly, VO_2_ max is considered a key physiological predictor of longevity in humans (29), further demonstrating that RE may have an important role in human healthspan/lifespan.

**Figure 4.**
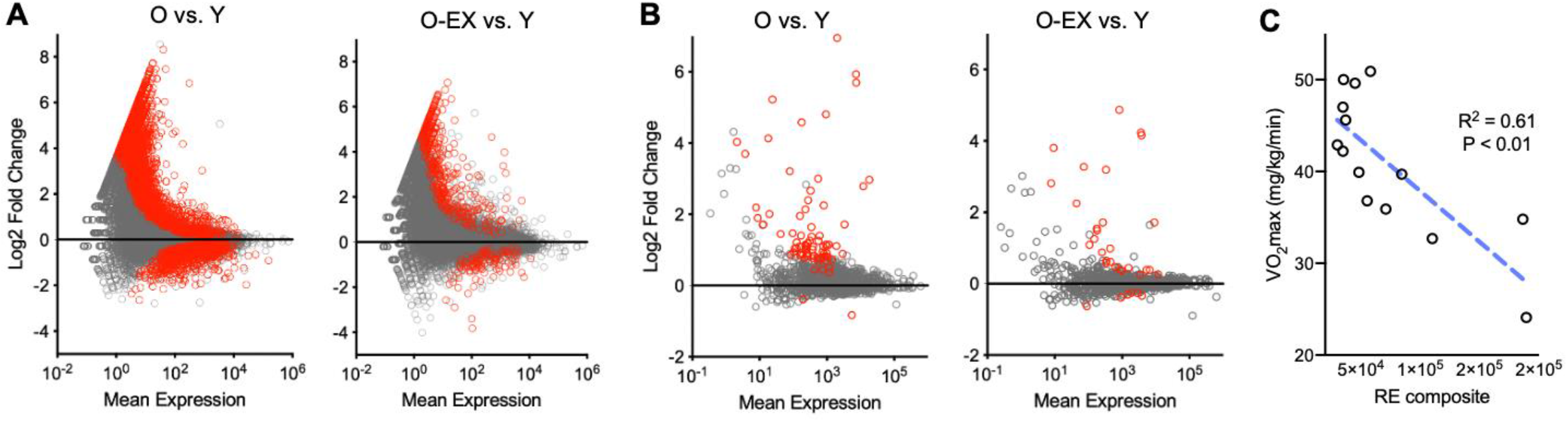
Long-term exercise is associated with reduced RE transcript expression in humans. **(A)** MA plots showing gene expression differences in peripheral blood mononuclear cells of older (O) vs. young (Y) sedentary, and older exercising (O-EX) vs. Y adults. N=4-5 matched samples per group. Note the ~2500 transcripts significantly increased/decreased with aging but largely reversed with exercise (red data points, FDR<0.1, identified using Deseq2). **(B)** MA plots showing RE transcript levels in the same samples/subjects. Note general upward shift and numerous significantly increased RE transcripts with aging (red, FDR<0.1, identified using Deseq2) that are largely reversed with exercise. **(C)** Correlation between maximal aerobic exercise capacity (VO_2_ max) and composite count of RE transcripts significantly increased with aging (O vs. Y) in all subjects. Human subjects characteristics in supplementary data file.

Collectively, our results support the growing idea that global RE dysregulation may be an important mechanism of aging (and not simply an adverse effect of the process). Reversing age-related RE transcript accumulation may be necessary for healthy aging, as our present findings show that health/lifespan-enhancing interventions consistently reduce RE expression. Indeed, we and others have reported age-associated increases in most types of RE transcripts (15, 16, 30), and this suggests a fundamental cellular mismanagement of RE with aging, which could have numerous deleterious effects. For example, our current study suggests that histone modifications, DNA damage and immune/inflammatory responses may be linked with RE dysregulation. Future investigations are needed to determine how and if these processes are specifically connected, as this could lead to novel strategies for additional, potentially complementary healthy aging interventions.

## MATERIALS AND METHODS

### RNA-seq datasets and availability

The data that support these findings can be found on the Gene Expression Omnibus under accession numbers (GEO): GSE92486 (long term CR), GSE131901 (8-month and 2-month CR and pharmacological treatments), GSE87565 (HFD), and GSE153100 (human exercise).

### Bioinformatics analyses

RE transcripts were quantified using TEtranscripts (31) and the RepEnrich2 algorithm (32, 33) as previously described (15), in order to confirm similar findings with RE different analysis platforms. Briefly, reads were trimmed, quality filtered with *fastp* (34), and then aligned to the genome (mm10 *Mus musculus* or Hg38 *Homo sapiens*) using Bowtie (RepEnrich) or the STAR aligner (TEtranscripts) (35). RE transcripts were then quantified using either TEtranscripts or RepEnrich, which are pipelines to quantify RE transcripts by individual total counts, class, and family. Gene expression counts were extracted from bam alignment files produced during TEtranscripts analyses, and differential expression analyses of both RE and genes were performed using Deseq2 software (36). WGCNA was performed according to standard procedures outlined by the analysis pipeline’s authors (37) using normalized gene and RE counts for all samples, and a minimum module size of 300 to capture broader groups of RE that correlated with sample traits (specific interventions). GO analyses of genes in the WGCNA modules were performed using the GOrilla algorithm (38), and specific GO modules were identified as terminal nodes in the directed acyclic graph produced by this program.

### Human subjects and RNA-seq samples

RNA-seq was performed on PBMC from twelve healthy young (18-22 years) and older (62-74 years) adults. Subjects were non-obese, non-smokers and healthy as assessed by medical history, physical examination, blood chemistries and exercise ECG, and small groups were selected to match characteristics as closely as possible. Young (n=5, 2 male) and older (n=5, 2 m) sedentary subjects performed no regular exercise (< 2 days/week, < 30 min/day), whereas older exercising subjects (n=4, 1 m) performed regular vigorous aerobic exercise (≥ 5 days/week, > 45 min/day) for the previous ≥ 5 years. The study conformed to the Declaration of Helsinki; all procedures were approved by the Institutional Review Board of the University of Colorado Boulder, and written informed consent was obtained from all subjects. Maximal oxygen consumption (VO_2_max) was assessed during treadmill exercise as previously described (39) and basic clinical measurements (e.g., blood pressure) were performed using standard techniques. PBMC were isolated from whole blood by traditional Ficoll gradient centrifugation, and RNA-seq and gene expression analyses were performed using standard methods as previously described (15, 40). Briefly, snap-frozen PBMC pellets were lysed in Trizol (Thermo), and RNA was recovered using a spin column kit (Direct-Zol, Zymo Research) that included a DNase I treatment to remove genomic DNA. Total RNA libraries were generated using Illumina Ribo-Zero kits to deplete ribosomal RNA, and libraries were sequenced on an Illumina NovaSeq 6000 platform to produce >40 M 150-bp single-end fastq reads per sample. Gene and RE expression analyses were performed as described above.

### Statistical Analyses

Differential expression of RE transcripts was quantified using the Deseq2 software as previously described using size factors to account for library size differences among samples (40). Chi-square analyses and heatmaps of increased/decreased RE and WGCNA modules were constructed using GraphPad Prism software, and Venn diagrams were generated using jvenn (41).

## Acknowledgments

This work is funded by the National Institute on Aging, National Institutes of Health (AG060302).

## Author contributions

D.W. designed the study, wrote the paper, generated and analyzed data, and provided conceptual insight; A.N.C. analyzed data, provided conceptual insight, and edited the paper; M.S., edited paper and provided conceptual insight; D.R.S. provided human PBMC samples, edited the paper, and provided conceptual insight; T.J.L. designed the study, wrote the paper, analyzed data and provided conceptual insight.

